# *p16.1* and *p16.2*, new HSPC markers, play redundant roles in zebrafish T-cell lymphopoiesis

**DOI:** 10.1101/2025.01.21.634058

**Authors:** Etienne Gomez, Roman A. Li, Julien Y. Bertrand

## Abstract

In the recent years, the zebrafish model has become a first-choice animal model in the field of hematopoiesis, due to extra-uterine development and optical transparency that allow an easy observation of early stages of hematopoietic stem and progenitor cells (HSPCs) development. Here, we characterized the function of two undescribed genes, *si:ch211-214p16.1* and *si:ch211-214p16.2* (shortened *p16.1* and *p16.2*) that are expressed by HSPCs, as early as their emergence. By combining different strategies, we have demonstrated that *p16.1* and *p16.2* play redundant roles in embryonic thymopoiesis. Knocking-down both genes at the same time – or one gene in the mutant background for the other gene – impaired HSPCs commitment towards lymphoid fate and homing to the thymus. However double mutant embryos did not show any phenotype. As these double-mutant animals were viable, their thymocytes were compared to their wild-type counterparts by RNA sequencing, revealing that double mutant thymocytes compensated the lack of *p16.1* and *p16.2* by upregulating genes involved in chemotaxis and cell migration. One of these transcripts, *ccr9b*, was indeed validated as an important regulator of thymic homing in double mutant embryos, only. In summary, we found two new markers of embryonic HSPCs, functionally linked to lymphoid fate. The description of such markers will be important for our understanding of heterogeneity among emerging HSPCs in the embryo.

## INTRODUCTION

Hematopoietic stem cells (HSCs) are responsible for the constant regeneration of blood lineages throughout life, therefore, they represent interesting tools for regenerative medicine against different kinds of genetic-related blood disorders. Indeed, many therapeutical approaches have already taken advantages of the accessibility of these cells in patients, as they can be harvested from the bloodstream after (McCredie, Hersh, and Freireich 1971; Anguita-Compagnon et al. 2010). Ex vivo, HSPCs can be modified using viruses (gene integration) or directly through genome editing (Ghosh, Thrasher, and Gaspar 2015; Naldini 2019; Ferrari et al. 2021) and then reinfused into patients to rescue the initial deficiencies. These therapeutical strategies involve high number of HSPCs, which are sometimes themselves malfunctional due to inflammatory stress (Sobrino et al. 2023). One strategy to circumvent these drawbacks is to directly derive induced HSPCs from a non-hematopoietic source, such as fibroblasts or human induced pluripotent cells. In order to achieve this goal, it is necessary to reproduce the molecular events that occur during embryogenesis in these cultured cells both at the cell and non-cell autonomous levels. This seems to have been successfully achieved, but always at low levels of efficiency (Pereira et al. 2013; Batta et al. 2014; Barcia Durán et al. 2018), which suggests that our knowledge of this process is still incomplete, although recent studies show promising results (Piau et al. 2023)

Developmental hematopoiesis occurs in successive waves culminating with the emergence of definitive long-term reconstituting HSCs (Elsaid et al. 2020), originating from hemogenic endothelial cells located in the floor of the aorta in all vertebrates (Bertrand et al. 2010; Kissa and Herbomel 2010; Boisset et al. 2010). These cells then migrate through several niches to achieve their maturation: the fetal liver then the bone marrow in mammals, or the caudal hematopoietic tissue (CHT), then the kidney marrow in zebrafish (Mahony and Bertrand 2019). Over the years, through the use of lineage tracing protocols, combined with sorting/staining techniques and the use of better in vitro culture and in vivo transplantation protocols, many groups have shown heterogeneity in the pool of hematopoietic progenitors produced from the hemogenic endothelium, showing different potential and different fates in zebrafish (Tian et al. 2017; Ghersi et al. 2023; Xia et al. 2023), but also in mouse models (Crisan and Dzierzak 2016). These progenitors are now referred to as hematopoietic stem and progenitor cells (HSPCs). In the mouse embryo, the sequential emergence of restricted lympho-myeloid progenitors can be monitored from the aorta, before the specification of true long-term reconstituting HSCs. This heterogeneity could be due to differential signals sent from the microenvironment, from an intrinsic heterogeneity of hemogenic endothelial cells, as shown in the zebrafish model (Ghersi et al. 2023; Xia et al. 2023), or from a combination of both events.

In our laboratory, we focus on genes of unknown function highly expressed in hemogenic endothelium and/or HSPCs, in order to find new pathways involved in the specification of HSPCs from the hemogenic endothelium, as well as to understand the molecular/genetic hierarchy governing HSPC heterogeneity. We used published single cell RNA sequencing datasets to find new targets, and in this study, we focus on two genes: *si:ch211-214p16.1* and *si:ch211-214p16.2*, both located in tandem on chromosome 17. These two genes were highly expressed in the cluster “Blood-unknown #21” in a single cell RNAseq performed on all cells from the zebrafish trunk at 30 hours post fertilization (hpf)(Metikala, Casie Chetty, and Sumanas 2021). This “Blood-unknown” cluster also expressed the highest levels of *myb*, *tal1* and *gfi1aa*, three genes related to emerging HSPCs. *si:ch211-214p16.2* was also recently associated with the pre-HSCs cluster in another single cell RNAseq dataset performed on endothelial cells (Sahai-Hernandez et al. 2023). And finally, we could find that these two genes were highly associated with two clusters: hemogenic endothelium and lymphoid-HSPC-2, using the online tool *Daniocell* (Sur et al. 2023).

Both genes are listed as “processed transcripts” with no protein product on the genome browser *Ensembl.* Yet for both genes, we found two possible open reading frames, coding for small proteins. We established the expression pattern of these two genes and show that both are expressed along the zebrafish aorta as early as 36 hpf. Their expression then shifts to the CHT and the thymus, at 48 and 96 hpf, respectively. In this study, we tried to elucidate the function of these two genes in HSC emergence and fate. By using a morpholino strategy, we find that these two genes are likely involved in either the commitment of early hematopoietic progenitors towards lymphoid fate or directly in their differentiation into *rag2*-positive thymocytes. Single mutants and even double-mutant embryos did not show any lymphoid phenotype, however, when performing transcriptome analysis on adult thymocytes from wild-type or double-mutant animals, we found that many genes involved in RhoGAP/GEF activity and migration were upregulated in double-mutant thymocytes, suggesting that these two new genes might be involved in this pathway. We provide here new markers to isolate lymphoid-biased HSPCs during zebrafish development.

## MATERIAL AND METHODS

### Ethical statement

Zebrafish were raised in accordance with FELASA and Swiss guidelines (Aleström et al. 2020) and all efforts were made to comply with the 3R guidelines.

### Zebrafish husbandry and transgenic lines

All zebrafish lines (AB* wild types, mutant lines and transgenic lines) were bred at 28.5°C in a 14:10 hours light:dark cycle. Embryos were obtained as previously described (Westerfield M. 2000). Embryo development was staged by hours post-fertilization (hpf) as previously described (Kimmel et al. 1995). The *Tg(ikfz1:GFP)*^fr101^ (referred to as *ikaros:eGFP*) was generated by the Boehm lab (Hess and Boehm 2012). The following mutant lines were also used: *cloche*^m39^ (Stainier et al. 1995), *mindbomb1*^ta52b^ (Itoh et al. 2003), *cmyb*^t25127^ (Soza-Ried et al. 2010).

### Morpholino purchase, storing and injections

Splice morpholinos specifically targeting *p16.1*, *p16.2* and *ccr9b* were purchased from Gene Tools (GeneTools, LLC, Oregon). While *ccr9b-MO* was already described (Punwani et al. 2016), we designed the following morpholinos: *p16.1-MO*, TGCCTTTGTCCTACACAAAATCAGA, and *p16.2-MO*, ATGAAAAGCAGTGCCTACCTCAGTT (Supplemental Figure S1A-B).Their efficiency was assessed by qPCR, using the following primers (see figure S1 C-D). p16.1-F-BamHI: AAGGATCCACCATGAAAATGAATCAAATTT, p16.1-R-XhoI: TTCTCGAGAGTTCAGTGCATGAAGC TGAGG, p16.2-F-BamHI: AAGGATCCACCATGAATAAATTATGGATTTT, p16.2-R-XhoI: TTCTC GAGTGTTCAGGATCTGCTGTGGATGA.

Lyophilized morpholinos were resuspended in 300μl of water for molecular biology (PanReac AppliChem) and stored at room temperature. Prior to injection, morpholinos stock solutions were heated for 10 minutes at 70°C. 1pl of morpholino solution was injected in one-cell stage embryos at the following concentrations: p16.1-MO = 2ng/nl, *p16.2-MO* = 8ng/nl. The *ccr9b-MO* was injected at the concentration of 4ng/nl as previously described (Punwani et al. 2016).

### Generation of the *p16.1^Δ4bp^, p16.1^Δ945bp^* and *p16.2^Δ464bp^*mutant alleles

Mutant alleles were generated using the CRISPR/Cas9 system for mutagenesis. The Cas9 enzyme was co-injected with gRNAs targeting *p16.1* or *p16.2* at the one-cell stage embryo. Adult fish from the cas9/gRNAs injected F0 presenting mutated *p16.1* or *p16.2* alleles in the germline were identified and mated to generate an F1 generation that would theoretically contain embryos somatically carrying mutated *p16.1* and *p16.2* alleles. By screening F1 fishes and mating them we managed to establish the following mutant lines: single mutant lines *p16.1*^Δ4/Δ4^ *; p16.2*^wt/wt^ and *p16.1*^wt/wt^ *; p16.2*^Δ464/Δ464^ and double the double mutant lines *p16.1* ^wt/Δ945^ *; p16.2*^Δ464/Δ464^ and *p16.1* ^Δ945/Δ945^ *; p16.2*^Δ464/Δ464^. gRNAs were synthesized from pDR274 vector using T7 transcriptase with the following oligonucleotides designed as their target sites: gRNA-16.2-F3, TAGGTCCAGAACTGAACCTTAC, gRNA-16.2-R3, AAACGTAAGGTTCAGTTCTGGA, gRNA-16.2-F5, TAGGCGAGCATCAATGTGGGAG, gRNA-16.2-R5, AAACCTCCCACATTGATGCTCG, gRNA-16.1-F2, TAGGTTAAAAGCAGCATCCCG, gRNA-16.1-R2, AAACCGGGATGCTGCTTTTAA, gRNA-16.1-F3, TAGGTGTTGTTTGTCGGTTGCG, gRNA-16.1-R3, AAACCGCAACCGACAAACAACA, TSE3-p16.1-oligo1, TAGGCCACCGATGCTGCAGGAC, TSE3-p16.1-oligo2, AAACGTCCTGCAGCATCGGTGG.

### Simple and fluorescent wholemount *in situ* hybridization

Wholemount *in situ* hybridization (WISH) were performed following the protocol as previously described (Thisse and Thisse 2008). Embryos were fixed overnight in a PBS solution containing 4% paraformaldehyde. Antisense RNA probes were generated using SP6/T7 RNA polymerase (Roche) with DIG or FITC labelled oligonucleotide mix (Roche). Simple WISH was performed using an anti-DIG antibody coupled with the alkaline phosphatase and signal was revealed by a metabolic reaction using Nitro-Blue Tetrazolium (NBT) and 5-Bromo-4-chloro-3-indolyl phosphate (BCIP) at room temperature. Fluorescent in situ hybridization was performed as previously described (Bertrand et al. 2007). Reactions were developed using anti-DIG-horseradish peroxidase (HRP) and anti-FITC-HRP antibodies (Roche). *gata-1*-FITC was detected with a mouse anti-FITC antibody (Roche). AlexaFluor-488 or -594 tyramide substrates (InVitrogen) were used according to manufacturer’s instructions to further amplify signals.

### Flow cytometry and cell sorting

Flow cytometry was performed on dissociated cell suspensions from dissected embryo or adult thymus. Embryo were chemically dissociated in PBS 0.9x /4% liberase (Sigma-Aldrich) at 33°C for 2h and resuspended in 0.9× PBS-1% fetal calf serum. Adult thymi were mechanically dissociated in PBS with 1% fetal calf serum. Dead cells were stained using SytoxRed (Molecular probes). Samples were analyzed on a BD LSR Fortessa and sorted with a BD FACS Aria Fusion. Cells were collected in 350μl RLT Plus lysis buffer from RNeasy Plus minikit for RNA extraction (QIAGEN) and stored at -80°C until RNA extraction and cDNA synthesis. FCS files were analyzed using FlowJo.

### RNA extraction and cDNA synthesis

RNA was extracted from cell lysate (sorted cell or whole embryo) in RLT Plus lysis buffer using the RNeasyPlus minikit for RNA extraction (QIAGEN). cDNA was synthetized from RNA solutions using the qScript cDNA SuperMix (QuantaBio).

### Quantitative real-time PCR

Quantitative real-time PCR (RT-qPCR) was performed on cDNA solutions using SensiFAST SYBR lo-ROX mix (Meridian biosciences) and run on a QuantStudio 3 machine by Thermofisher with the primers listed in the table under *morpholino purchase, storing and injections*.

### Imaging and quantification

WISH against *ikaros* and *rag1* were imaged in glycerol using Olympus MVX10 microscope at a 2.5 times magnification. Living fluorescent samples were imaged on a glass slide using an Olympus IX83 microscope at 10 times magnification. All pictures were acquired under the TIFF format using the CellSens Dimension software (Olympus). *ikaros* and *rag1* signal areas were measured manually using the Fiji ImageJ software and are expressed in arbitrary units.

## RESULTS

### *p16.1* and *p16.2* are both expressed by HSPCs throughout embryonic development

Both genes present a similar structure, consisting of two exons and one intron, and contain putative open reading frames (ORFs) that would encode a 77-residue peptide for *p16.1* and a 68-residue peptide for *p16.2.* In order to define the anatomical and temporal frame of *p16.1* and *p16.2* expression during zebrafish embryogenesis, we performed wholemount *in situ* hybridization (WISH) against the two genes using DIG-labelled antisense riboprobes. Expression of both genes was detected by WISH from 34hpf up to 5dpf by HSPCs in the different hematopoietic organs of embryonic life (figure 1A-C). At 2dpf, expression of *p16.1* and *p16.2* was detected at the site of emergence of HSPCs, the dorsal aorta (figure 1A). At 3.5dpf, the expression of both *p16.1* and *p16.2* was lost along the dorsal aorta, but both genes were now expressed in the CHT as well as in the thymus (figure 1B). Eventually, at 5dpf, *p16.1* and *p16.2* expression was found in all hematopoietic sites (figure 1C): the CHT, the thymus, along the aortic region and in the kidney glomeruli, which will ultimately become the definitive hematopoietic niche in adult zebrafish. At earlier stages, we could detect *p16.1* and *p16.2* by fluorescent ISH, and show that they were specifically expressed by cells lining the floor of the aorta, presumably hemogenic endothelial cells (figure 1D). As expected, the expression of both *p16.1* and *p16.2* was completely lost in *cloche* and *mindbomb* mutant embryos (figure 1E), reinforcing our hypothesis that these two genes mark HSPCs, and are expressed by vascular cells, probably downstream of *Notch* signaling. These two genes were also lost in *cmyb* mutants (figure 1E), showing that their expression tightly relies on hematopoietic fate. The expression pattern of these two genes appears to reflect their allocation to HSPC clusters in different single cell RNA sequencing data sets. Therefore, we decided to further investigate the functions of these two genes in developmental hematopoiesis.

**Figure 1.**
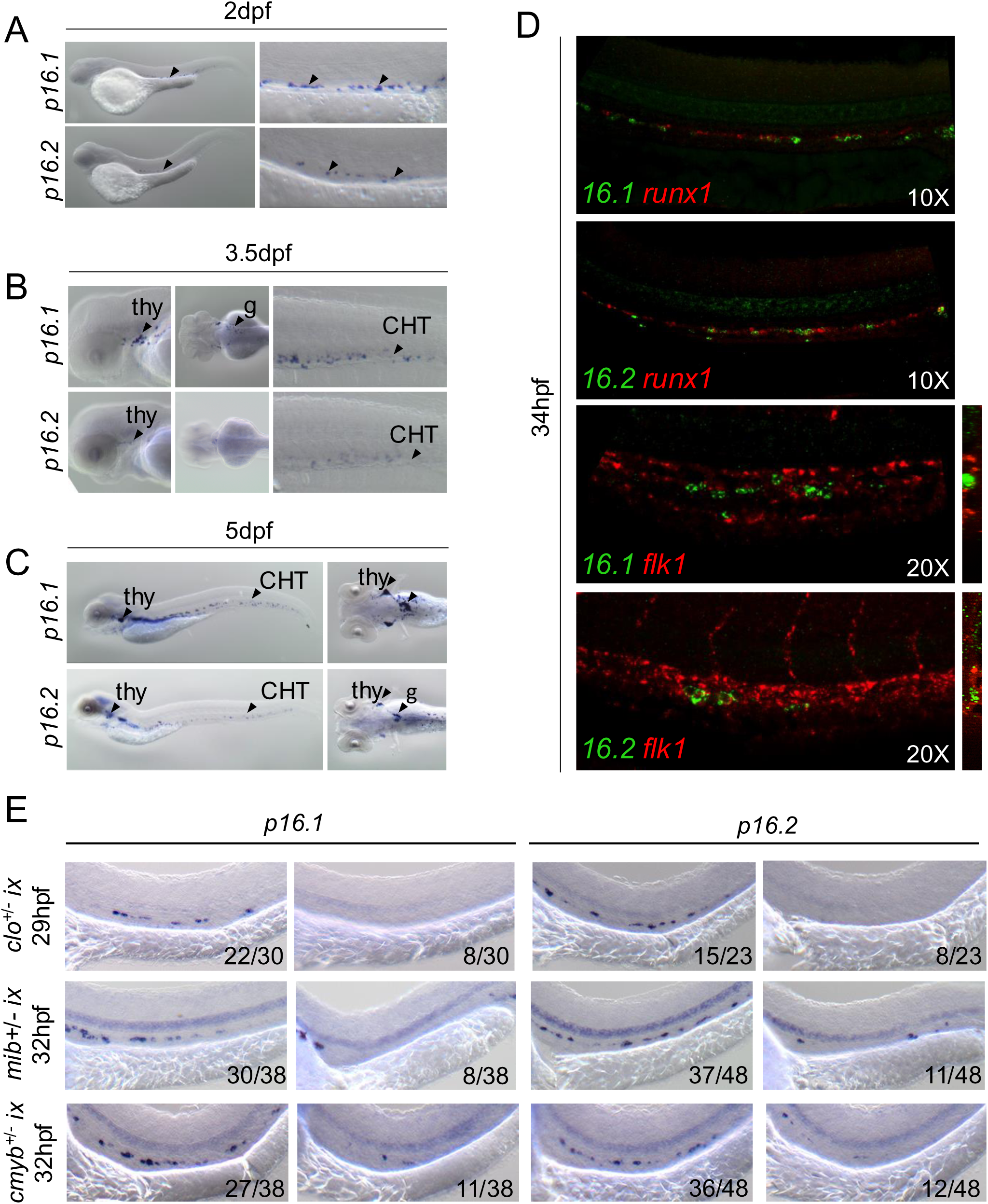
Expression pattern of *p16.1* and *p16.2* in the zebrafish embryo. (A-C) WISH against *p16.1* and *p16.2* showing expression of both genes in hematopoietic sites of the embryo. (A) At 2dpf *p16.1* and *p16.2* are detected in the dorsal aorta (black arrows). (B) At 3.5dpf, *p16.1* and *p16.2* in the thymus (thy) in the CHT. *p16.1* is detected in the kidney glomeruli (g). (C) At 5dpf, *p16.1* and *p16.1* are detected in the thymus (thy), the CHT and in the kidney glomeruli (g). (D) fluorescent HIS of showing expression of *p16.1* and*p16.2* in the ventral wall of the dorsal aorta at 34hpf. (E) WISH against *p16.1* and *p16.2* in various mutant lines, *cloche*, *mindbomb* and *cmyb* mutants showing a loss of *p16.1* and *p16.2* expression *clo*^-/-^*, mib*^-/-^ and *cmyb*^-/-^ embryos.

### *p16.1* and *p16.2* play redundant roles in embryonic T-lymphopoiesis

In order to investigate the function(s) of *p16.1* and *p16.2* in embryonic hematopoiesis, we combined two parallel strategies, using morpholinos (supplemental figure S1) and CRISPR/Cas9 mutant lines, derived in our laboratory (supplemental figure S2). We then performed WISH against hematopoietic markers upon transient knock-down/loss of both genes. Therefore, we injected two splice morpholinos, MO-p16.1 and MO-p16.2, that target *p16.1* and *p16.2*, respectively to get double morphants (DMO). A non-specific control morpholino (MO-ctrl) was injected as a negative control (figure 2A). Double knock-down of *p16.1* and *p16.2* did not seem to affect neither *runx1* nor *cmyb* expression pattern at 24hpf and 60hpf, respectively (data not shown). However, DMO embryos presented a reduction of the T-cell marker *rag1* in the thymus at 5dpf as compared with age matched control embryos (figure 2B). To confirm the specificity of the morpholino treatment on the observed phenotype, we decided to phenocopy our morpholino with mutant animals. We generated two distinct *p16.1* and *p16.2* mutants (supplemental figure 2A-C). The *p16.1*^Δ4^ mutant allele consist of a 4bp deletion that leads to a frame shift in the ORF of the gene and to a premature stop codon (supplemental figure S3). The *p16.2*^Δ464^ allele consists of a 464bp deletion that encompasses the ORF of the gene (supplemental figure S4). Of note, none of the single mutants showed any hematopoietic phenotype, when we checked *runx1*, *cmyb* or *rag1* at 28hpf, 36hpf and 5dpf, respectively (data not shown). We then performed a single MO-*p16.2* knock-down in *p16.1*^-/-^ mutant embryos (and a *p16.1* knock-down in *p16.2*^-/-^ embryos) to try to recapitulate the effect of a double knock-down of *p16.1* and *p16.2.* Interestingly, single knock-downs of *p16.1* and *p16.2* in the opposite corresponding knock-out line led to a decrease of *rag1* at 5dpf, similar to what we observed in DMO embryos (figure 2D-G). Altogether, this data allows us to conclude that *p16.1* and *p16.2* play a redundant role, not during HSPC emergence from the hemogenic endothelium, but rather in T-cell lymphopoiesis during developmental hematopoiesis.

**Figure 2.**
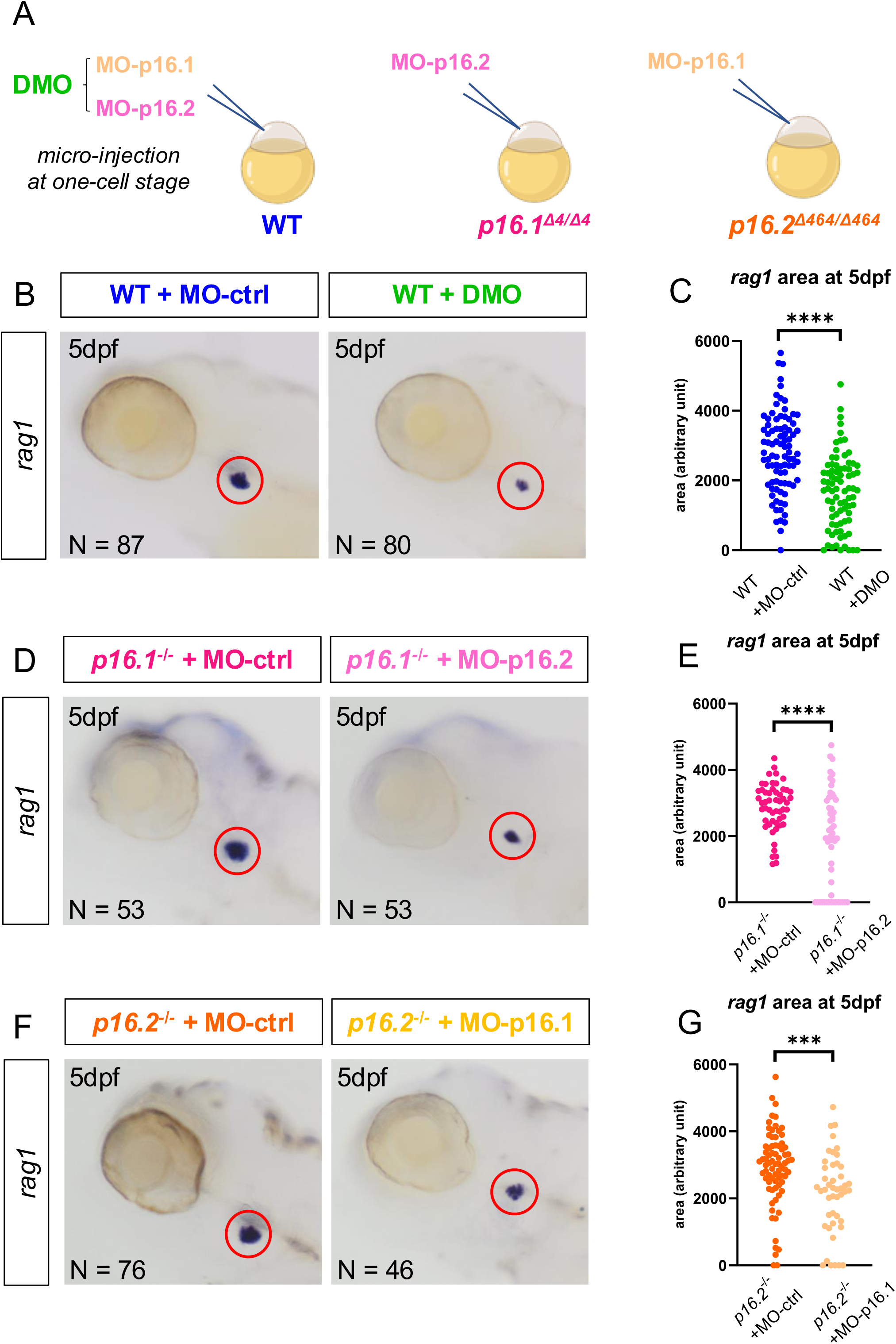
*p16.1* and *p16.2* knock-down in wild-type and single mutant lines. (A) Injection of MO-p16.1 and MO-p16.2 (DMO) in wild type embryos, or MO-p16.2 in single mutants *p16.1*^Δ4/Δ4^, or MO-p16.1 in single mutants *p16.2*^Δ464/Δ464^. (B) WISH against *rag1* at 5dpf in wild-type embryos injected with MO-ctrl (left panel) or DMO (right panel). (C) Quantification of the *rag1* signal area in arbitrary unit (a.u.) at 5dpf in wild-type embryos injected with MO-ctrl (N=87, mean=2790, SD=1144.4) and DMO (N=80, mean=1685, SD=1050.8). p<0.0001. (D) WISH against *rag1* at 5dpf in *p16.1*^-/-^ embryos injected with MO-ctrl (left panel) or MO-p16.2 (right panel). (E) Quantification of the *rag1* signal area in arbitrary unit (a.u.) at 5dpf in *p16.1*^-/-^ embryos injected with MO-ctrl (N=53, mean=2873, SD=727.04) and MO-p16.2 (N=53, mean=1887, SD=1480.2). p<0.0001. (F) WISH against *rag1* at 5dpf in in *p16.2*^-/-^ embryos injected with MO-ctrl (left panel) and MO-p16.1 (right panel). (G) Quantification of the *rag1* signal area in arbitrary unit (a.u.) at 5dpf in *p16.2*^-/-^ embryos injected with MO-ctrl (N=56, mean=3084, SD=1117.1) and MO-p16.1 (N=46, mean=2139, SD=1187.5). p<0.0001. Significance is calculated with unpaired T-test and denoted as *p<0.05, **p<0.01, ***p<0.001, ****p<0.00001 and *ns* when the difference is not significant.

### *p16.1* and *p16.2* are involved in lymphoid commitment and thymic homing of HSPCs

The *ikaros:eGFP* fluorescent reporter line can be used as a marker of lymphoid progenitors in the CHT and in the thymus (Hess and Boehm 2012; Mahony, Pasche, and Bertrand 2018). We therefore used this line to perform double knock-down of *p16.1/.2* and analyzed DMO embryos by cytometry and fluorescent microscopy to measure lymphoid progenitors in the CHT at 48hpf, as well as thymic colonization between 48hpf and 72hpf (figure 3A). By cytometry, we could determine that DMOs harbored a decrease of the *ikaros:eGFP*^high^ cell population in the CHT at 48hpf (figure 3B-C, Supplemental figure S5), which corresponds to a loss of lymphoid-committed progenitors (Mahony, Pasche, and Bertrand 2018). Interestingly, this decrease of the lymphoid precursors in the CHT at 48hpf resulted in a delay of thymic colonization, when monitored by fluorescent microscopy (figure 3D-E). At 72hpf, when 100% of MO-ctrl injected embryos had their thymus colonized by at least one *ikaros:eGFP*^high^ cell, 25% of DMO embryos still had no thymic colonization (figure 3D-E). At 72hpf, we separated embryos into three distinct groups: MO-ctrl embryos (MO-ctrl), DMO embryos with colonized thymi (DMO-c) and DMO embryos with non-colonized thymi (DMO-nc). These three groups of embryos were then fixed separately at 4dpf and WISH against the endogenous *ikaros* gene was performed (figure 3F). Measurement of the *ikaros* signal showed that there was no significant difference between MO-ctrl and DMO-c but that there was a significant *ikaros* decrease in DMO-nc embryos (figure 3F-G). The data strongly suggests that the decrease of *rag1* observed at 5dpf in DMOs likely results from either a defect in lymphoid commitment in the CHT, or a problem in thymic homing, or a combination of both.

**Figure 3.**
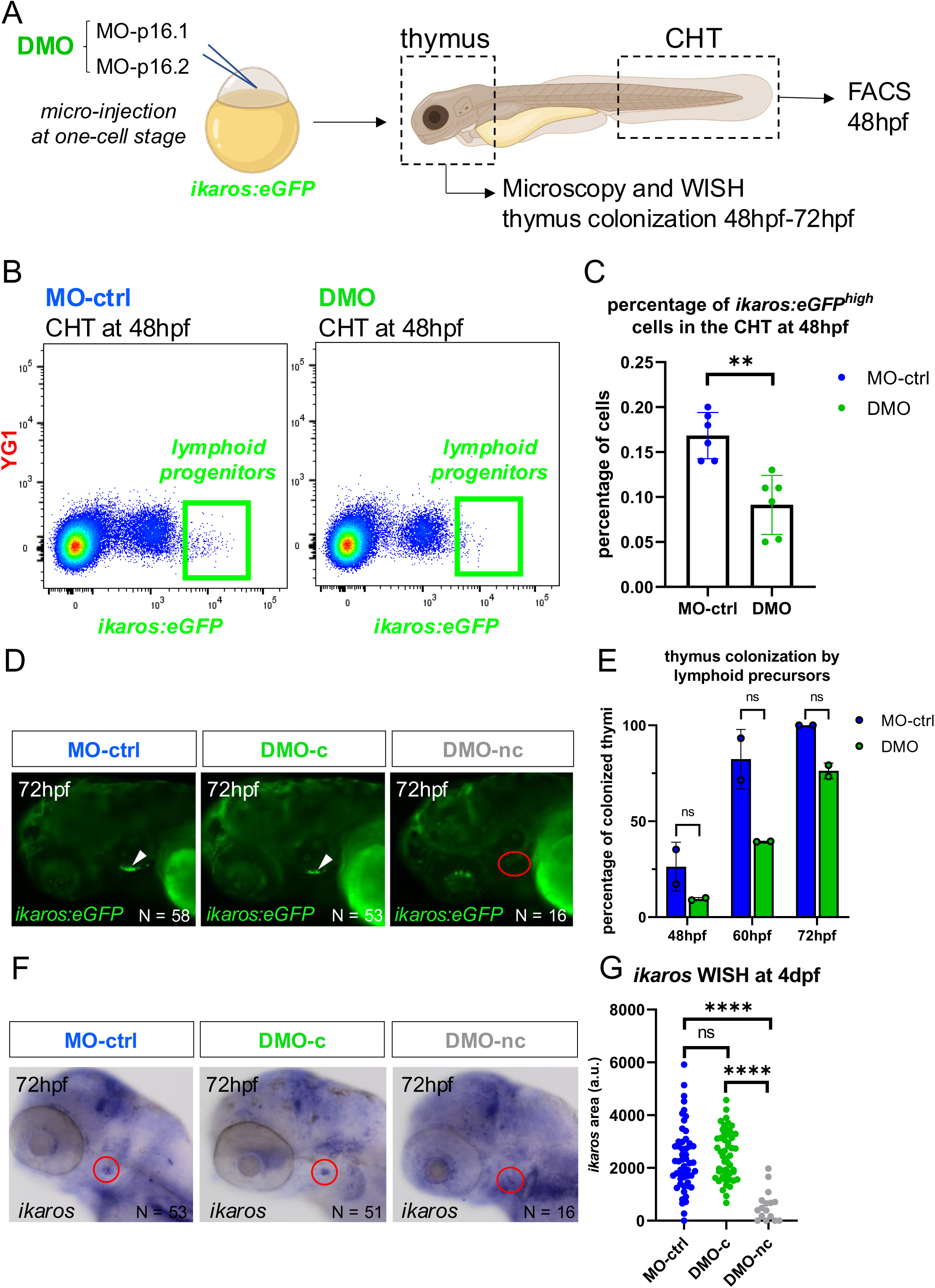
*p16.1* and *p16.2* knock-down lead a to a decrease of lymphoid precursors in the CHT and a delayed thymus colonization. (A) Experimental approach to evaluate the effect of *p16.1* and *p16.2* double knock-down on lymphoid progenitors by injecting MO-p16.1 and MO-p16.2 (DMO) into the *ikaros:eGFP* fluorescent reporter line. (B) FACS plots showing a decrease of the *ikaros:eGFP*^high^ population (green gate) corresponding to lymphoid progenitors in the CHT at 48hpf in DMO (right panel) as compared with MO-ctrl injected embryos (left panel). (C) Graph showing the frequency of *ikaros:eGFP*^high^ cells (lymphoid progenitors) in the CHT at 48hpf in MO-ctrl (N=6, mean=0.1683, SD=0.0256) and DMO (N=6, mean=0.0913, SD=0.0328) injected embryos. p=0.0011. (D) Fluorescent microscopy showing the state of thymus colonization by lymphoid precursors at 72hpf: 100% of MO-ctrl injected embryos (n=2, N=58) present colonized thymi, whereas DMO injected embryos can be segregated into two groups DMO-c (73.25% of DMO, n=2, N=58) with colonized thymi and DMO-nc (23.75% of DMO, n=2, N=16) with non-colonized thymi. (E) Graph showing the percentage of thymi colonized by at least one lymphoid precursor at 48hpf, 60hpf and 72hpf in MO-ctrl and DMO injected embryos (n=2, mean_48hpf MO-ctrl_ 26.27%, SD_48hpf MO-ctrl_=12.77, mean_48hpf DMO_=9.44%, SD_48hpf DMO_=0.79, mean_60hpf MO-ctrl_=82.36% SD_60hpf MO-ctrl_=15.49, mean_60hpf DMO_=39.41%, SD_60hpf DMO_=0.18 mean_72hpf MO-ctrl_=100%, SD_72hpf MO-ctrl_=0, mean_72hpf DMO_ =76.25%, SD_72hpf DMO_=4,12. (F) WISH against *ikaros* at 4dpf on MO-ctrl, DMO-c and DMO-nc (n=2). (G) Quantification of the *ikaros* signal is in arbitrary unit (a.u.) at 4dpf in MO-ctrl (N=53, mean=2348, SD=1231.388), DMO-c (N=51, mean=2470, SD=916.961) and DMO-nc (N=16, mean=571, SD=588.938). Significance is calculated with unpaired T-test and denoted as *p<0.05, **p<0.01, ***p<0.001, ****p<0.00001 and *ns* when the difference is not significant.

### *p16.1*^-/-^*p16.2^-/-^* double mutant embryos present comparable T-cell lymphopoiesis as compared with *p16.1^+/-^p16.2^-/-^*siblings

Morpholinos are very convenient tools to explore the effects of a loss of function of genes. However, their effect is only transient. We therefore turned to double knock-out animals, but as *p16.1* and *p16.2* are positioned so close on chromosome 17, it was impossible to generate double mutant embryos by crossing our single mutant parents. We decided to generate a double mutant line using CRISPR/Cas9 directed to the *p16.1* locus in the *p16.2*^-/-^ background, which generated a new *p16.1*^Δ945^ mutation (figure 4A, supplemental figure S6). The resulting fish, *p16.1*^+/-^*p16.2*^-/-^, will be referred to as HET. We had to incross these animals to get double mutant embryos, *p16.1*^-/-^*p16.2*^-/-^, referred to as double-mutants or DMU. Both HETs and DMUs animals are viable, grow into adulthood and are fertile. Due to the short intergenic distance between *p16.1* and *p16.2* the two loci are transmitted as one block: therefore, crossing HET fish with DMU fish leads to a 50/50% distribution of HET and DMU genotypes in the progeny (figure 4B). Since our data suggests that *p16.1* and *p16.2* play a redundant function in T-cell lymphopoiesis, we hypothesized that HET embryos would present a T-cell lymphopoiesis comparable to wild type embryos and we then investigated hematopoiesis in DMU embryos. We performed WISH against *ikaros* and *rag1* at 4.5dpf and 5dpf, respectively, on embryos resulting from a DMUxHET breeding, but could not observe any difference in thymic lymphopoiesis between HETs and DMUs (figure 4C-D), suggesting that DMU embryos completed T-cell lymphopoiesis just as well as HET embryos. To confirm that earlier stages of hematopoiesis showed normal progression in DMU embryos, we in-crossed heterozygous double mutants *p16.1*^+/^ *p16.2*^+/-^ fish and performed WISH against *runx1* and *cmyb* at 28hpf and 72hpf, respectively. Indeed, we found that DMU embryos (*p16.1*^-/-^*p16.2*^-/-^) showed similar hematopoiesis as compared to their double heterozygous (*p16.1*^+/-^*p16.2*^+/-^) and wild types (*p16.1*^+/+^*p16.2*^+/+^) siblings (supplemental figure S7). As the phenotype of double knock-out was very different from our transient double knock-down, we sought to understand the molecular pathway involved in a potential compensatory mechanism.

**Figure 4.**
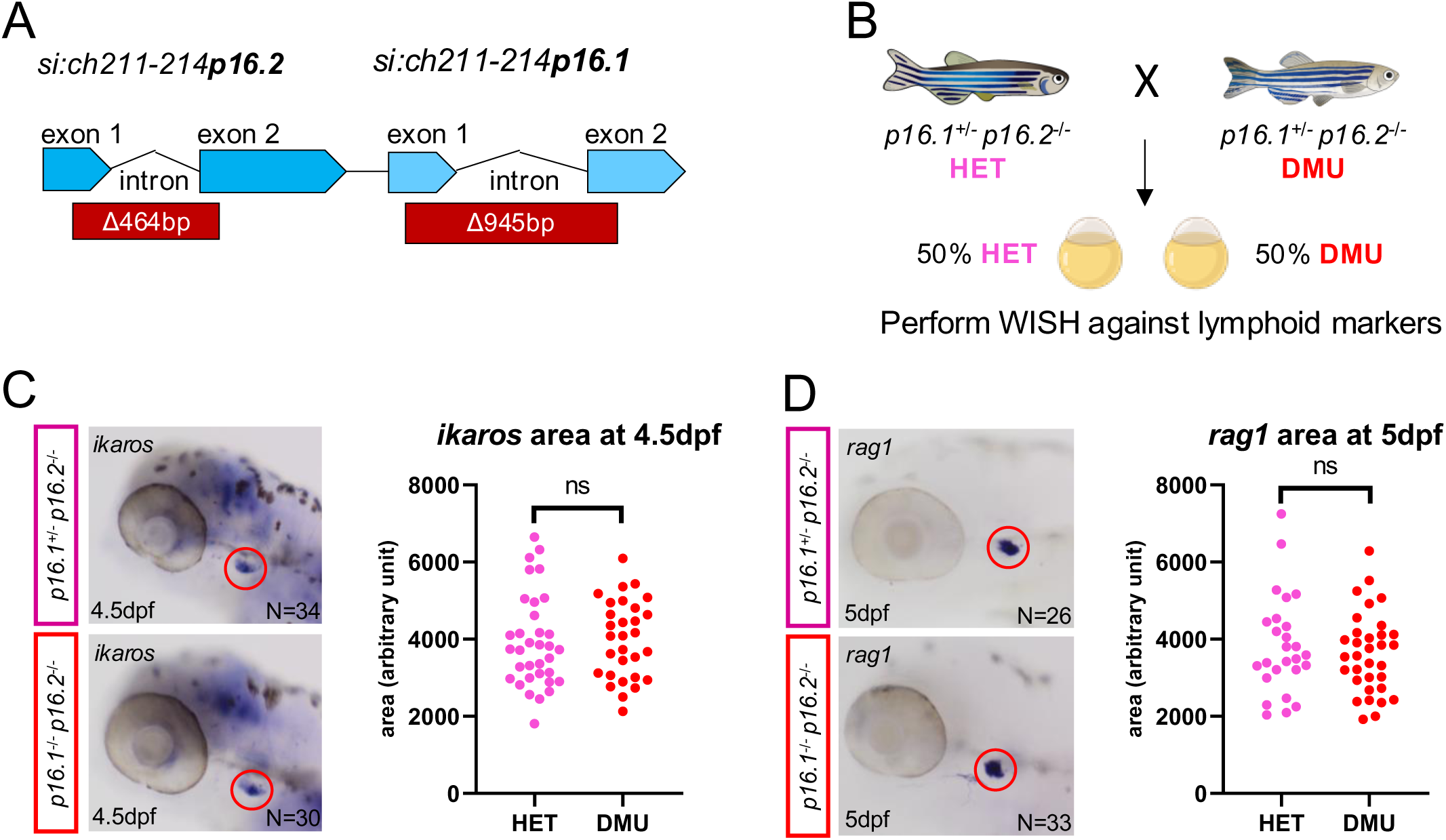
Characterization of embryonic thymopoiesis in double mutants. (A) Schematic view of the *p16.1*^Δ945^ and the *16.2*^Δ464^ alleles in HET (*p16.1*^+/-^;*16.2* ^-/-^) and DMU (*p16.1*^-/-^;*16.2* ^-/-^). (B) Experimental approach to characterize thymopoiesis in double mutants by crossing HET with DMU fish. (C) WISH against *ikaros* at 4.5dpf in HET and DMU siblings (left panel) and quantification of *ikaros* signal area in arbitrary unit (a.u.) (right panel). N_HET_=34, mean_HET_=3927, SD_HET_=1197.359 N_DMU_=30, mean_DMU_=4000, SD_DMU_=1008.426. n=3, p=0.7932. (D) WISH against *rag1* at 5dpf in HET and DMU siblings (left panel) and quantification of *rag1* signal area in arbitrary unit (a.u.) (right panel). N_HET_=26, mean_HET_=3827, SD_HET_=1278.306, N_DMU_=33, mean_DMU_=3632, SD_DMU_=1042.051. n=3, p=0.5216. Significance is calculated with unpaired T-test and denoted as *p<0.05, **p<0.01, ***p<0.001, ****p<0.00001 and *ns* when the difference is not significant.

### Adult DMU thymocytes differ from their wild-type counterparts

The zebrafish model has been shown to be very capable to by-pass deleterious effects of mutations by modifying the transcriptomic landscape of their cells (El-Brolosy et al. 2019). In order to investigate potential compensatory modifications in T-cell precursors in DMU embryos, we sorted *ikaros:eGFP+* cells from the thymus of adult WT and DMU fish and performed bulk RNA sequencing (figure 5, supplemental figure S8). First, we noticed that the profiles of GFP expression were different between both genotypes. While WT animals show a homogeneous population of *ikaros:GFP*-positive cells, DMU animals showed a bi-modal expression of GFP (figure 5A). When we compared both transcriptomes, bioinformatic analysis showed that 1580 genes were differentially expressed, among which 574 and 1006 were upregulated and down-regulated in DMU thymocytes, respectively. When we subjected this list to GO Term analysis, we noticed that most of the differentially expressed genes belonged to pathways involved in the immune response, and also to chemotaxis and cell migration (supplemental figure S9). We observed that DMU thymocytes expressed fewer transcripts involved in T cell commitment, such as *notch1b*, *irf4a*, *il2rb*, *il7r*, as well as less *p16.1* (figure 5B). *irf4a* was previously associated to lymphoid commitment in the CHT (Huang et al. 2019), which is impaired in double morphants (figure 3B-C). Finally, a lot of genes associated with cytotoxic responses were also down regulated, such as genes coding for perforins and granzyme (figure 5B). Noticeably, *rag1* and *cd247* (coding for the CD3-associated zeta chain) were up-regulated in DMU cells (figure 5B). Cell migration is mediated by a number of surface receptors, most of them G-protein coupled receptors, as well as by the intracellular Rho pathway, controlled by RhoGAPs and RhoGEFs, which controls numerous aspects of migration, such as cytoskeleton (Nayak et al. 2013). DMU cells expressed higher levels of *ccr9b*, an orthologue to mammalian CCR9, a receptor for CCL25, expressed by the thymic stroma, as well as higher levels of *cxcr4b* and *ackr3a* (also known as *cxcr7a*), both receptors for *Cxcl12* (figure 5C). These 3 genes code for GPCR that are involved in HSPC migration towards the hematopoietic niche (Jenkinson et al. 2007; Calderón and Boehm 2011). DMU cells also expressed higher levels of direct or indirect activators of the Rho pathway (*fam13a, arhgef9a, abr, rac1l, rap1gap2b*) (figure 5D). In particular, *rac1l*, not expressed by WT thymocytes and highly expressed by DMU cells, is a paralog of *rac2* that is important for T-cell progenitors migration to the thymus in the mouse and the zebrafish (Faroudi et al. 2010; X. Lu et al. 2020). Conversely, DMU cells expressed lower levels of *ripor2*, which codes for a negative regulator of RhoA and negatively controls T cell migration (Megrelis et al. 2018), and lower levels of *s1pr5a*, which was recently involved in the retention of T cells in circulation (Evrard et al. 2022) (figure 5C-D). Overall, it seems that DMU animals can bypass *p16.1* and *p16.2* to establish their T cell compartment, by increasing the levels of genes involved in thymic colonization in hematopoietic progenitors.

**Figure 5.**
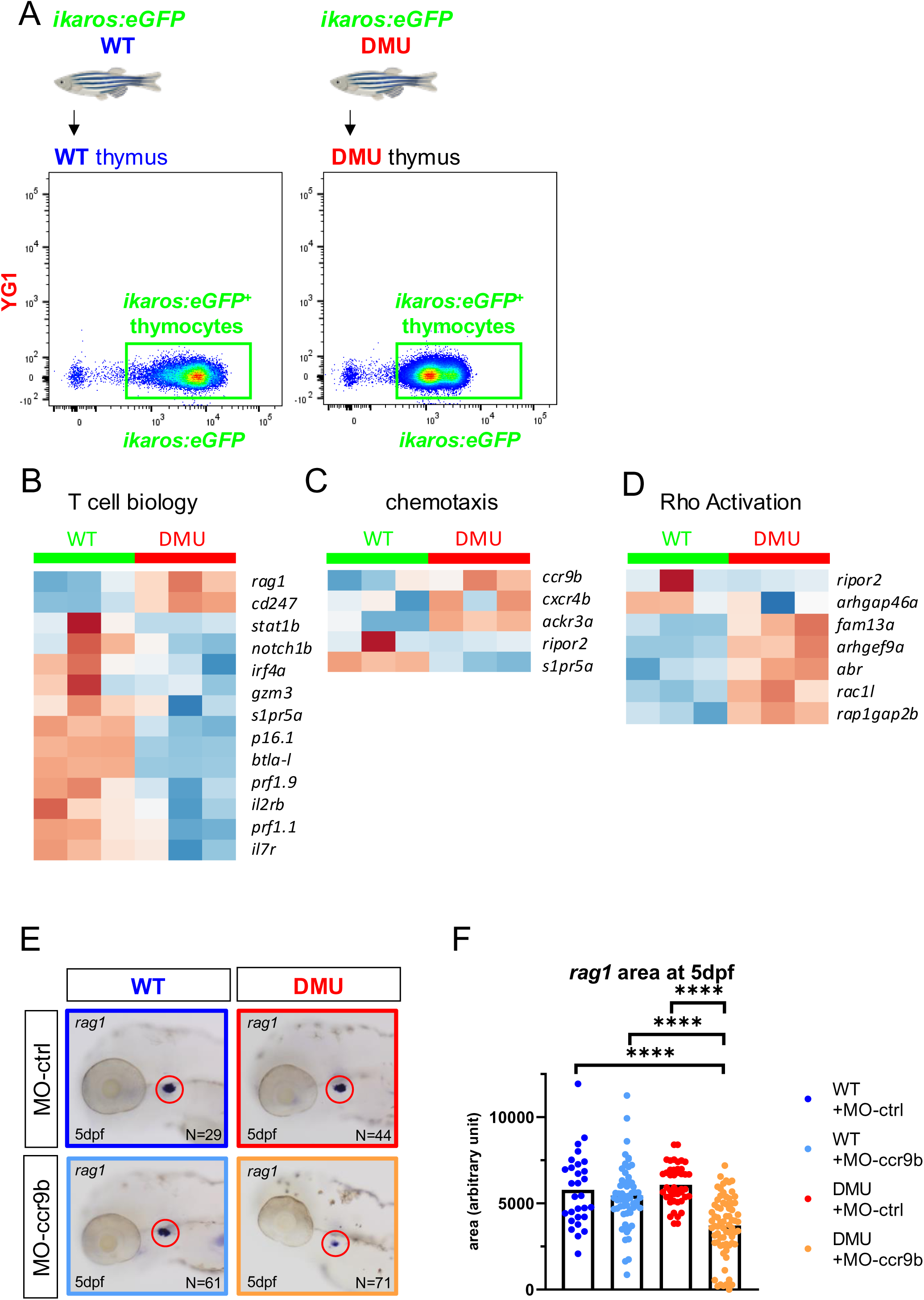
Bulk RNAseq on T-cell progenitors from DMU fish reveals upregulation of several genes involved in thymopoiesis. (A) Sorting T-cell and T-cell progenitors (*ikaros:eGFP*^+^) from thymi of adult WT and DMU fish.(B-D) Heatmaps showing differentially expressed genes between WT and DMU lymphoid progenitors. These genes are involved in chemotaxis, Rho-GAP/GEFs and T-cell function. Volcano plot showing differentially expressed genes between WT and DMU T-cell progenitors. Upregulated transcripts of *ccr9b*, *abr*, *arhgef9a* and *racl1* genes known to be involved in thymopoiesis. (E) WISH against *rag1* at 5dpf in WT and DMU injected with MO-ccr9b or MO-ctrl showing a decrease of *rag1* signal in DMU upon *ccr9b* knock-down. (F) Quantification of *rag1* area in arbitrary unit (a.u.) in WT+MO (N=29, mean=5634, SD=2105.066) WT+MO-ccr9b (N=61, mean=5452, SD=1776.531), DMU+MO-ctrl (N=44, mean=6079, SD=1164.346). DMU+MO-ccr9b (N=71, mean=3711, SD=1737.374) n=3. Significance is calculated with unpaired T-test and denoted as *p<0.05, **p<0.01, ***p<0.001, ****p<0.00001. For clarity, statistically non-significant differences are not notified.

### DMU HSPCs rely on *ccr9b* to colonize the thymus and differentiate into *rag2***^+^** thymocytes

Among the genes upregulated in DMU thymocytes, we found *ccr9b*, a paralog of *ccr9a,* and both genes are orthologues of mammalian *CCR9. CCR9* encodes a G-protein coupled receptor that specifically binds CCL25, a chemokine expressed by thymic epithelial cells, and necessary for hematopoietic progenitors to colonize the thymus (M.-A. Wurbel et al. 2000; M. A. Wurbel et al. 2001; Liu et al. 2006). In zebrafish, *ccr9a* plays a similar role. It is expressed by HSPCs as early as the CHT stage (Wang et al. 2015), while *ccl25a* and *ccl25b* are expressed by the thymic anlage (I.-N. Lu et al. 2012; Hess and Boehm 2012; Wang et al. 2015) and disrupting the *ccr9a/ccl25a* axis delays and prevents HSPCs homing to the thymus (Huang et al. 2019), whereas *ccr9b* does not seem involved in this process (Punwani et al. 2016), as it is not expressed by HSPCs at these stages (Xia et al. 2021). Therefore, we hypothesized that completion of lymphopoiesis in DMU embryos may at least partially rely on the upregulation of *ccr9b* and that transient knock-down of *ccr9b* using a *ccr9b*-specific splice morpholino should impact thymopoiesis as observed in DMOs. Therefore, we injected *ccr9b*-MO in WT and DMU embryos and performed WISH against *rag1* at 5dpf. Consistent with our hypothesis, only DMU embryos presented a reduced *rag1* area at 5dpf, whereas WT embryos were unaffected by the knock-down of *ccr9b* (figure 5E-F). From this, we can conclude that *ccr9b* is only necessary to achieve normal thymopoiesis in DMU embryo, as a compensatory mechanism due to the loss of *p16.1* and *p16.2*.

## DISCUSSION

We are constantly looking for new functional markers of embryonic hematopoietic progenitors, in order to: 1- establish new tools to purify embryonic HSPCs, 2- understand the molecular pathways involved in definitive HSPC emergence, and 3- decipher HSPC heterogeneity, as increasing evidence show that many flavors of HSPCs arise from the hemogenic endothelium (or from multiple distinct hemogenic endothelia?) in the zebrafish embryo, as well as in mammals. In this study, we focused on two new genes of unknown function*: si:ch211-214p16.1* and *si:ch211-214p16.2*, referred to as *p16.1* and *p16.2*. Both genes frequently appear in the top enriched genes associated to HSPCs in different single-cell RNA sequencing data sets, but no function has been associated with these new markers. Through analysis of their expression patterns, we validated that these two new genes are associated with the emergence of definitive HSPCs. Indeed, these two genes are first expressed along the floor of the aorta, after HSPC specification, and their expression pattern then shifts to the CHT and the thymus, similarly to HSPCs (Murayama et al. 2006; Kissa et al. 2008). Concerning their expression in adult hematopoiesis, single-cell RNA sequencing show that these genes are also associated with the T/NK lineages as well as with HSCs/Blood progenitors to a lesser extend (Tang et al. 2017). Therefore, the evidence pointed to a potential role in thymopoiesis during embryogenesis.

We tested this hypothesis by combining approaches using both morpholinos and the use of mutant lines that we develop with CRISPR/Cas9 tools. While single morphants and single mutants did not show any hematopoietic phenotype, we observed that double morphants showed a strong delay in the onset of thymopoiesis, which likely originated from a decrease in lymphoid commitment from newly generated HSPCs at the level of the CHT. This suggested that both *p16.1* and *p16.2* were necessary for normal thymopoiesis. This was further consolidated by the use of the *p16.1*-MO in *p16.2* mutants, and vice versa, showing that both genes are redundant in this process. However, hematopoiesis was completely normal in double mutants. The deletions we generated with CRISPR/Cas9 encompass most of the genes, so we excluded genetic compensation as a mechanism to rescue these double mutant embryos, as this process has been shown to occur as a consequence of a null mutation, involving a premature termination codon (El-Brolosy et al. 2019). However, HSPC apparently committed normally towards lymphoid fate. To understand how they could circumvent the absence of *p16.1* and *p16.2*, we analyzed the transcriptome of thymic *ikaros:GFP* cells, sorted from adult double mutants, which are viable, and compared them to wild-type thymic control cells (also *ikaros:GFP*-positive). Although thymopoiesis seemed unaffected in these double mutants, we could actually show that a number of genes were deregulated between controls and double mutants. Among them, we found a high number of genes involved in RhoGAP/GEF regulation such as *abr* and *arhgef9a*. These proteins regulate the activity of Rho and Rac/Cdc42 proteins, which are involved in numerous biological processes, such as cell migration (Faroudi et al. 2010) and even T cell activation downstream of TCR signaling (Sanui et al. 2003; Singleton et al. 2009). The migration aspect retained our attention as double morphants show a defect in thymus colonization. Moreover, among the deregulated genes in thymocytes sorted from adult double mutants, we found that the expression of *ccr9b* was abnormally elevated, compared to controls. *ccr9a* and *ccr9b* are both zebrafish paralogs of the CCR9 chemokine receptor in mammals, which is expressed by thymic-seeding hematopoietic progenitors. CCR9-positive hematopoietic progenitors indeed follow the gradient of CCL25, expressed by the thymic epithelium, and colonize the thymus to differentiate into mature T cells (M. A. Wurbel et al. 2001; Liu et al. 2006). In the zebrafish model, *ccr9a* is believed to be the main driver of thymic colonization, as 1-thymopoiesis cannot occur in the absence of *ccr9a* (Wang et al. 2015), and as 2- *ccr9b* is basically not expressed by embryonic HSPCs (Xia et al. 2021), and therefore not involved in this initial thymic colonization. We therefore extrapolated our adult transcriptomic results, and hypothesized that we could have similar results in embryonic HSPCs, and tested the role of *ccr9b* in thymic colonization. While the morpholino targeting *ccr9b* did not affect thymopoiesis in wild-type embryos, it did somehow decreased thymopoiesis in double mutant embryos. It is therefore likely that, similarly to their adult counterparts, double mutant embryonic HSPCs upregulate *ccr9b*. Ultimately, it appears that *p16.1* and *p16.2* may either play a role directly in the molecular pathway that drives cell migration (Rho GAP/GEF, CCR9-CCL25 axis) or somehow contribute to the genetic regulation of these coding genes. At this stage, we cannot conclude precisely on the role of these two new genes, and being unable to find an orthologous gene in mammals does not help. The status of these two genes is still under consideration on ensemble.org, but our own results show that these two genes could eventually code for small proteins of 77 or 68 amino-acids, for *p16.1* and *p16.2*, respectively. Although a comprehensive annotation of short open reading frames has been performed in the zebrafish genome (Pauli, Valen, and Schier 2015) it only starts to emerge in human genomics (Leong et al. 2022), and hopefully functional equivalents for *p16.1* and/or *p16.2* will be annotated at some point. Having more tools will allow us to find molecular partners, and eventually lead to better understanding the function of these two genes.

Nonetheless, we now have a valuable tool to mark embryonic HSPCs in the zebrafish embryo, and more precisely, lymphoid-biased HSPCs. Indeed, the expression of *p16.1* and *p16.2* seems to be tightly linked to lymphoid fate decision, as their expression becomes restricted to thymocytes during early embryogenesis, and since their expression is also highly enriched in adult lymphocytes (unpublished results from E. Gomez). Different groups have now proposed that the hemogenic endothelium either gives rise to different types of HSPCs, endowed with different differentiation capabilities, or that the heterogeneity would already be encrypted at the level of the hemogenic endothelium itself (Ghersi et al. 2023; Xia et al. 2023). The presence of these two lymphoid-related genes, *p16.1* and *p16.2*, could therefore mark the first T-cell biased HSC-independent progenitors, as they were originally described by Tian and colleagues (Tian et al. 2017). Further studies, using new tools to perform cell-lineage tracing, will be necessary to decipher the molecular and cellular mechanisms underlying this functional heterogeneity. This will only be made possible with the characterization of new functional markers.

## Acknowledgements

We would like to acknowledge all past and current members from the Bertrand lab for their comments and suggestions during this project. This work was originally supported by a Giorgi-Cavaglieri chair to JYB, and then by the swiss national foundation to JYB (310030_184814).

## Data availability

All raw data will be available on Yareta, following this link (will be delivered after acceptance of the manuscript). The accession number for the bulk RNA-sequencing data reported in this paper is Gene Expression Omnibus (GEO): GSExxxxxxx. (TBD)

## Author contributions

R.L. initiated the p16.1 project, established expression patterns for both genes and also generated the single CRISPR/Cas9 mutants. E.V.G. performed all other experiments. R.L., E.V.G. and J.Y.B. designed experiments. E.V.G. wrote the first draft of the manuscript, and all authors contributed to the final version.

## Disclosure and competing interest statement

The authors declare that they have no conflict of interest.

## Notes

### Competing Interest Statement

The authors have declared no competing interest.

