## supplementary data for "*p16.1* and *p16.2*, new HSPC markers, play redundant roles in zebrafish T-cell lymphopoiesis"

**Supplemental Figure S1**

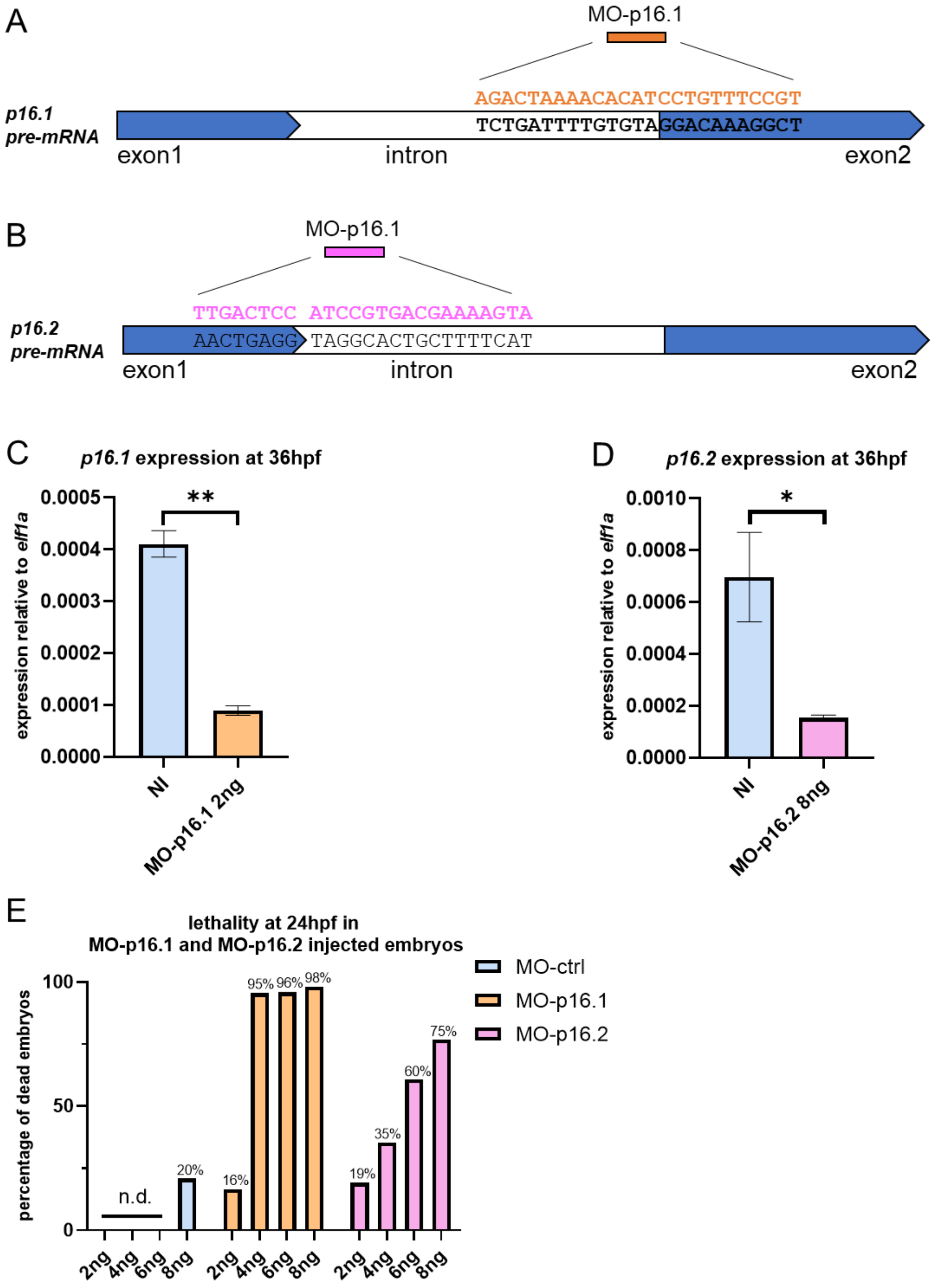

**Supplemental figure S1. Transient knock-down of *p16.1* and *p16.2* by splice morpholinos.** (A-B) schematic view of binding sites of (A) MO-p16.1 and (B) MO-p16.2 splice morpholinos at the intron-exon junction of *p16.1* and *p16.2.* (C-D) relative expression of *p16.1* (left panel) and *p16.2* (right panel) at 36hpf in non-injected (NI) and *p16.1* single morphants (MO-p16.1 2ng) or *p16.2* single morphants (MO-p16.2 8ng). (C) relative expression of *p16.1* at 36hpf in non-injected (NI) embryos (n=3, N = 3, $mean_{NI}$=0.00041 , $SD_{NI}$=2.25E-5) and in *p16.1* single morphants (n=3, N=3, $mean_{MO\text{-}p16.2\ 2ng}$=8.93E-5, $SD_{MO\text{-}p16.2\ 2ng}$=9.02E-6). p=0.0017. (D) relative expression of *p16.2* at 36hpf in non-injected (NI) embryos (n=3, N= 3, $mean_{NI}$=0.00069, $SD_{NI}$=0.00017) and in *p16.2* single morphants (n=3, N=3, $mean_{MO\text{-}p16.2\ 2ng}$=0.00015, $SD_{MO\text{-}p16.2\ 2ng}$=1.09E-5). p=0.0359. (E) Graph representing lethality at 24hpf in *p16.1* morphants, *p16.2* morphants and control embryos in a dose dependent manner (n=1, n.d.=not done).

**Supplemental Figure S2**

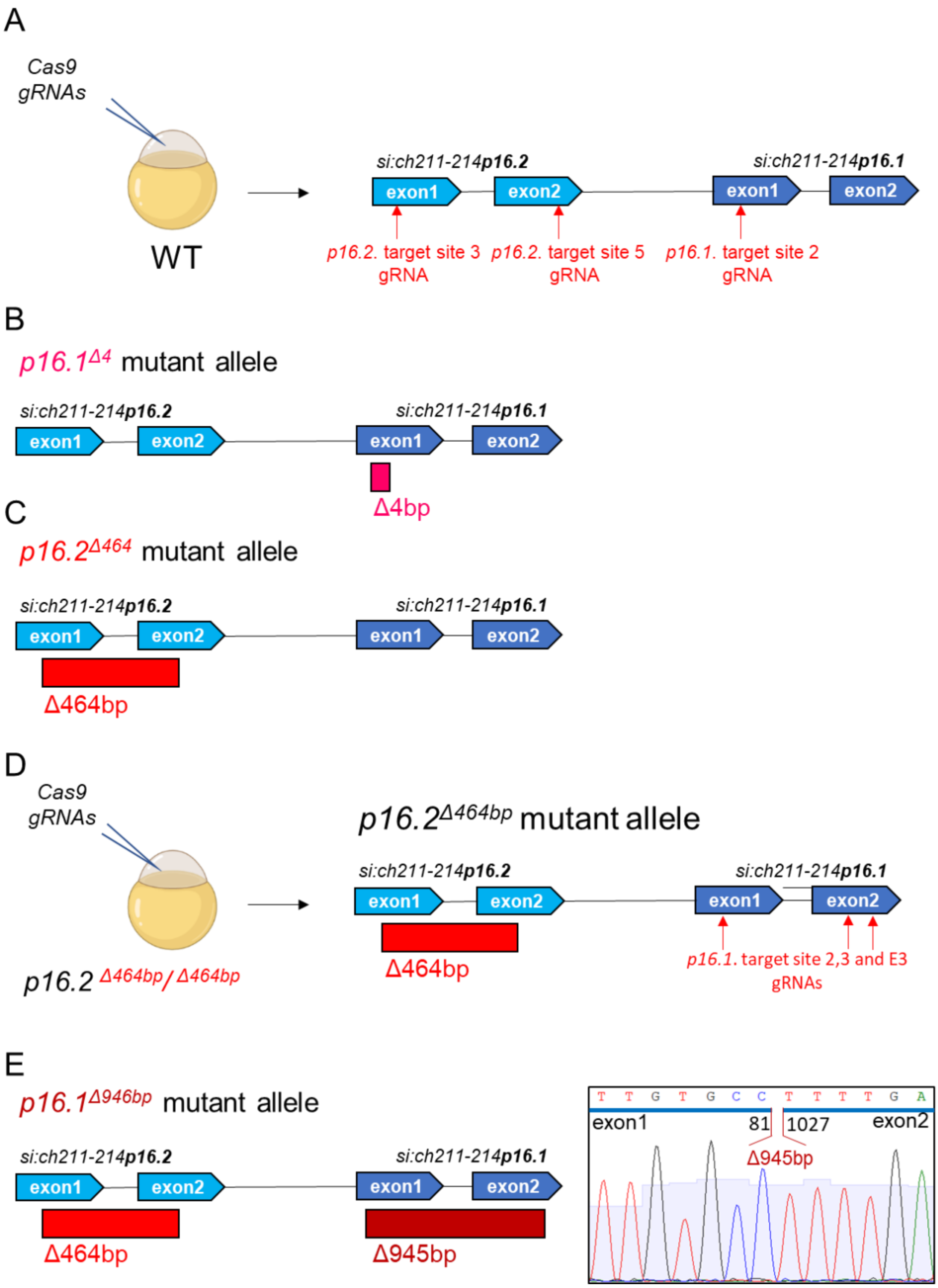

**Supplemental figure S2. Generation of the *p16.1$^{Δ4}$*, *p16.2$^{Δ464}$* and *p16.1$^{Δ945}$* mutant alleles using CRISPR/Cas9.** (A) Generation of single mutant lines by injecting Cas9 with gRNAs targeted against *p16.1* or *p16.2* in a wild type background. (B) The *p16.1$^{Δ4}$* allele consists of a 4bp deletion that induces a frameshift in the putative ORF of *p16.1* and leads to the formation of a premature STOP codon. (C) the *p16.2 $^{Δ464}$* allele consists of a 464bp deletion that encompasses the entire putative ORF of *p16.2*. (D) Generation of the double mutant line by injecting Cas9 with gRNAs targeted against *p16.1* in a *p16.2$^{Δ464/Δ464}$* mutant background. (E) The *p16.1$^{Δ945}$* mutation consists of a 945bp deletion.

### Supplemental Figure S3

#### *si:ch211-214p16.1* genomic sequence

CGACTACACATTTTCTGCCCCACTGAAAACATCTTGACAAGAGACCACCATGAAAATGAATCAAATTTTGAC

ATTGTGCCTCGGGATGCTGCTTTTAACCTTACCCTTTTCAACCTCAGAGgtaggcagtgttattttcaaact

cattaatctttgatttttttttttattcgtatttaagaaactttgatattttccttattacctgaatatga

atgcaaaacaaattattgcctccagttcttttaaaccaagtaattagaagtttgggtcatttgttaaaatgg

actcttggctagttcaccaaaactgaaaattcggccattaatcactcatcctccacttgttccaaatcaatt

tcttttttttcagttgaACACAAAAGAAGATATACTGAAGAATGTTGAGAAAAACAGCCATTATCTTCCTTTT

AATAATATTTTTCCGACTATGGATATCAATGGCTGCAGGTTTCCAACATTCTTCAGAACATTTAGTTTTGGG

CTCAACAGATAAAAAATCATAATGGTTTTAGTGTGAGTAAATGGTGAGGACATATTTGGGTGAATGACTGAT

GAAGCATCATAAAGAATTACAAATGTAATAACAATAAAGTGCATTTTAAGGCACCTTCTATCAAAATAACTT

TAATTCAACTTTAAATAACTTTAAGCTGATTTCACATTGCATATGAAAAAGGCGGGTTACCTAAAACAAAAC

AAAAGAACAAATGAAAAAAATTTTTTAATTACTTTATAAAAGCCAGATGTCCGTTAGGGTATACAACTGCAC

ATTTTTCACAATAGTGGACATTATAATGATCTGCCTGTTTATTCTGATTTTGTGTAGGACAAAGGCTCAGCA

AAAGATGGTGAGCAAAAACCAGCCTCGCAACCGACAAACAACACCAATACCAACACTGCGGGGGCCACCGAT

GCTGCAGGACTGGATGTGTTGCCCACCATGCTGTGTTTACTGATTCCAGCTGCCACCCTCAGCTTCATGCAC

TGAACTGGCTGGAGAGTGTTTTGATGTAACCCAGAAAGACGGACAGCCAGTCACATGAAGAGAGAGATTTTC

CACAGTTCTGGAAGACACATAAATAAAGCAAGATGCCACAGCAATTCAAATTAGAAAAAGATTTTGTTTATT

TGTTAATATAAATATTTCTGGAAAAAAAAAAAACGGCTTTAAAAGGACAACACCCAACAAAGCCACACAGAT

GGACTCTTGCCCCATTTTTGAAAATACCCATCAGCTGTGATTTCCTTGTTCAAAATTAAACATGATAATTCA

GTTATACAATATTTTTGGCTTGATTGTAATTATAATGATTAAGAATATGTCTATATGCACAATTAAAGTGGA

TGAT

**Legend** :

EXONS (bases in capital letters)

intron (bases in lowercase)

gRNA TARGET SITES

**DELETED BASES in the *p16.1*$^{\Delta 4bp}$ mutated allele**

ATG start codon of the ORF

TGA stop codon of the ORF

**Supplemental figure S3. Genomic DNA sequence of *si:ch211-214p16.1* showing the location of the deletion in the *p16.1*$^{\Delta 4}$ allele.** The *p16.1*$^{\Delta 4}$ allele consists of a 4bp deletion that induces a frameshift in the putative ORF of the gene.

### Supplemental Figure S4

#### *si:ch211-214p16.2* genomic sequence

ACACTCACCCTCGGTCCAGAACTGAACCTTACTGGAAGAACACAACACAGCATTAACCATGAATAAATTATG
GATTTTGTGTTTCGTGGTGCTCATCATGAGCAAACCATTTTCAACTGAggtaggcactgcttttcattgtta
tctcattgagttttagaagtaaaaggaatcatttgtttcactcatttgcaaactgtgttcatattttacata
ttttaactagaatgtgtagatctacaaacagctccagaatgtttttattttttattttatctgtgatgctgt
aaagtgactgcttcatgtctgatgtagGAGAGTCAGAAGTCAAGTGAAAACACTGTCCCATCGCCCAGCGCA
TCTACTAACTCCAGTGCTGTAGGACTGAGTCCTGTGCCCAAAACACTGTACCTTCTGTTCCCGGCTGCCACC
CTGGGCTTCATCCACAGCAGATCCTGAACATTCACCAAAGGCGAGCATCAATGTGGGAGTGGCAGAATAGAT
AGTCATTGACAGACAGAGCACAAATCTGCAAGACAGAACACAGCACAGCACAGCTCAAAGGATTTTTAACTA
ATGAAAATAAGAACTCCTCCAAGAAGGATTGTCTGGTTACGCTCAGTGGCAACTTCTCTTGGAAGAGCCCCT
TTCTTCTATTTGCTTCATGTGAATAAAGTTTAGCAGGGTAAACAACTCCACTACTTAAATCATGCGTTTGAA
AGATTCTATTTCTCTAATGGCATTATGTGTATTAATGTAATTGACTGGCAAGTGTGTCTGTATGCATGATTA
AACTGCTAAATGCC

**Legend** :

EXONS (bases in capital letters)

intron (bases in lowercase)

gRNA TARGET SITES

**DELETED BASES in the *p16.1$^{\Delta 464bp}$* mutated allele**

ATG start codon of the ORF

TGA stop codon of the ORF

**Supplemental figure S4. Genomic DNA sequence of *si:ch211-214p16.2* showing the location of the deletion in the *p16.2$^{\Delta 464}$* allele.** The *p16.1$^{\Delta 644}$* allele consists of a 464bp deletion that encompasses the putative ORF of the gene.

**Supplemental Figure S5**

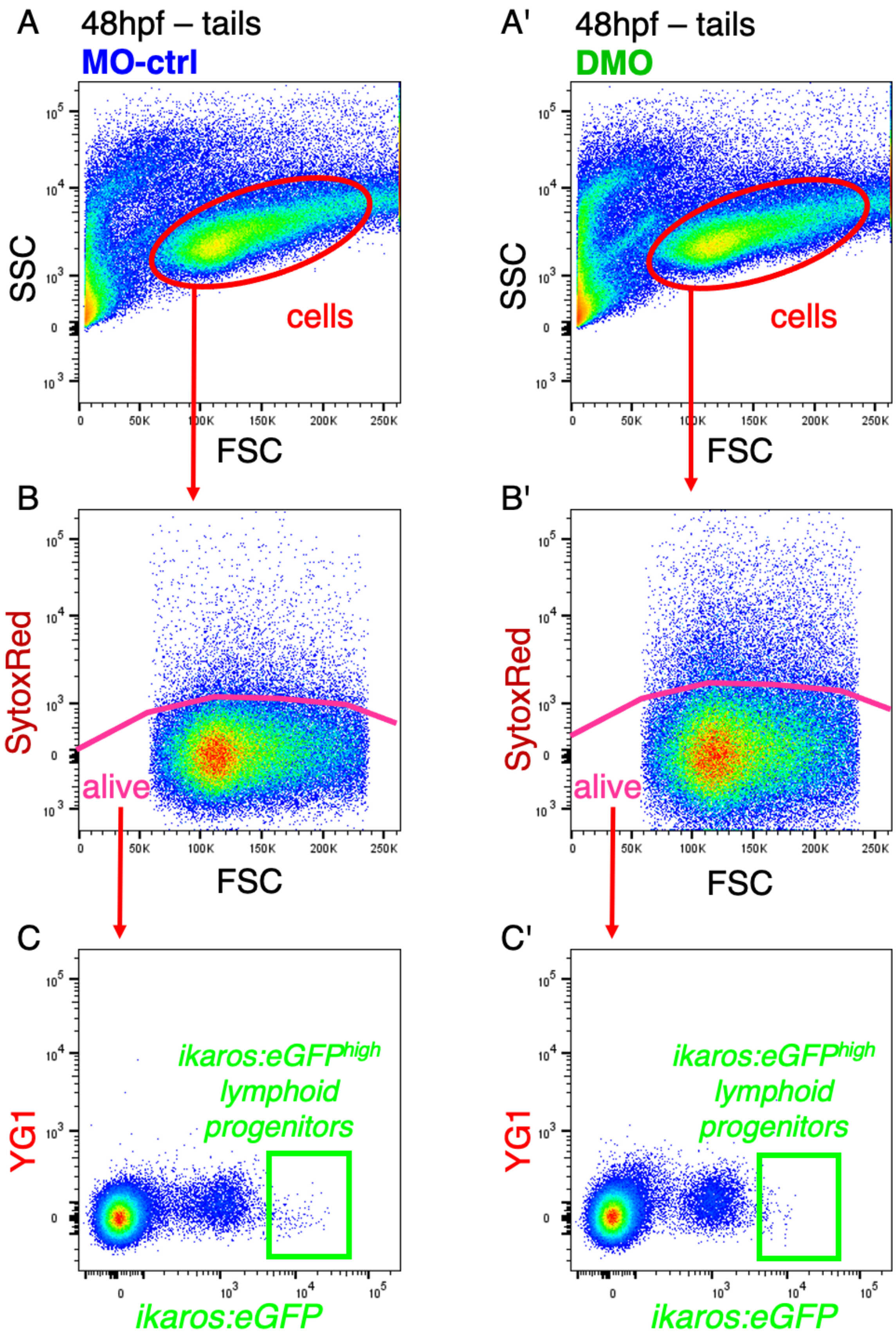

**Supplemental figure S5. Gating strategy to analyze lymphoid progenitors in the CHT by flow cytometry.** (A and A') Gating on the cells of interest (cells) within dissected tails based on the FSC/SSC parameters. (B and B') Selection of living SytoxRed$^{-}$ cells (alive) within cells of interest. (C and C') *ikaros:eGFP*$^{high}$ correspond to lymphoid progenitors in the CHT and present a lower relative abundance in DMO embryos as compared with MO-ctrl embryos.

**Supplemental Figure S6**

***si:ch211-214p16.1* genomic sequence**

CGACTACACATTTTCTGCCCCACTGAAAACATCTTGACAAGAGACCACCATGAAAATGAATCAAATTTTGAC

ATTGTGCCTCGGGATGCTGCTTTTAACCTTACCCTTTTCAACCTCAGAGgtaggcagtgttattttcaaact

cattaatctttgatttttttttttttattcgtatttaagaaactttgatattttccttattacctgaatatga

atgcaaaacaaattattgcctccagttcttttaaaccaagtaattagaagtttgggtcatttgttaaaatgg

actcttggctagttcaccaaaactgaaaattcggccattaatcactcatcctccacttgttccaaatcaatt

tctttttttcagttgaACACAAAAGAAGATATACTGAAGAATGTTGAGAAAAACAGCCATTATCTTCCTTTT

AATAATATTTTTCCGACTATGGATATCAATGGCTGCAGGTTTCCAACATTCTTCAGAACATTTAGTTTTGGG

CTCAACAGATAAAAAATCATAATGGTTTTAGTGTGAGTAAATGGTGAGGACATATTTGGGTGAATGACTGAT

GAAGCATCATAAAGAATTACAAATGTAATAACAATAAAGTGCATTTTAAGGCACCTTCTATCAAAATAACTT

TAATTCAACTTTAAATAACTTTAAGCTGATTTCACATTGCATATGAAAAAGGCGGGTTACCTAAAACAAAAC

AAAAGAACAAATGAAAAAAATTTTTTAATTACTTTATAAAAGCCAGATGTCCGTTAGGGTATACAACTGCAC

ATTTTTCACAATAGTGGACATTATAATGATCTGCCTGTTTATTCTGATTTTGTGTAGGACAAAGGCTCAGCA

AAAGATGGTGAGCAAAAACCAGCCTCGCAACCGACAAACAACACCAATACCAACACTGCGGGGGCCACCGAT

GCTGCAGGACTGGATGTGTTGCCCACCATGCTGTGTTTACTGATTCCAGCTGCCACCCTCAGCTTCATGCAC

TGAACTGGCTGGAGAGTGTTTTGATGTAACCCAGAAAGACGGACAGCCAGTCACATGAAGAGAGAGATTTTC

CACAGTTCTGGAAGACACATAAATAAAGCAAGATGCCACAGCAATTCAAATTAGAAAAAGATTTTGTTTATT

TGTTAATATAAATATTTCTGGAAAAAAAAAAAAACGGCTTTAAAAGGACAACACCCAACAAAGCCACACAGAT

GGACTCTTGCCCCATTTTTGAAAATACCCATCAGCTGTGATTTCCTTGTTCAAAATTAAACATGATAATTCA

GTTATACAATATTTTTGGCTTGATTGTAATTATAATGATTAAGAATATGTCTATATGCACAATTAAAGTGGA

TGAT

**Legend** :

EXONS (bases in capital letters)

Introns (bases in lowercase)

gRNA TARGET SITES

**DELETED BASES in the *p16.1*$^{Δ945}$ mutated allele**

ATG start codon of the ORF

TGA stop codon of the ORF

**Supplemental figure S6. Genomic DNA sequence of *si:ch211-214p16.1* showing the location of the deletion in the *p16.1*$^{Δ945}$ allele.** The *p16.1*$^{Δ4}$ allele consists of a 945bp deletion that removes most of the putative ORF of the gene.

**Supplemental Figure S7**

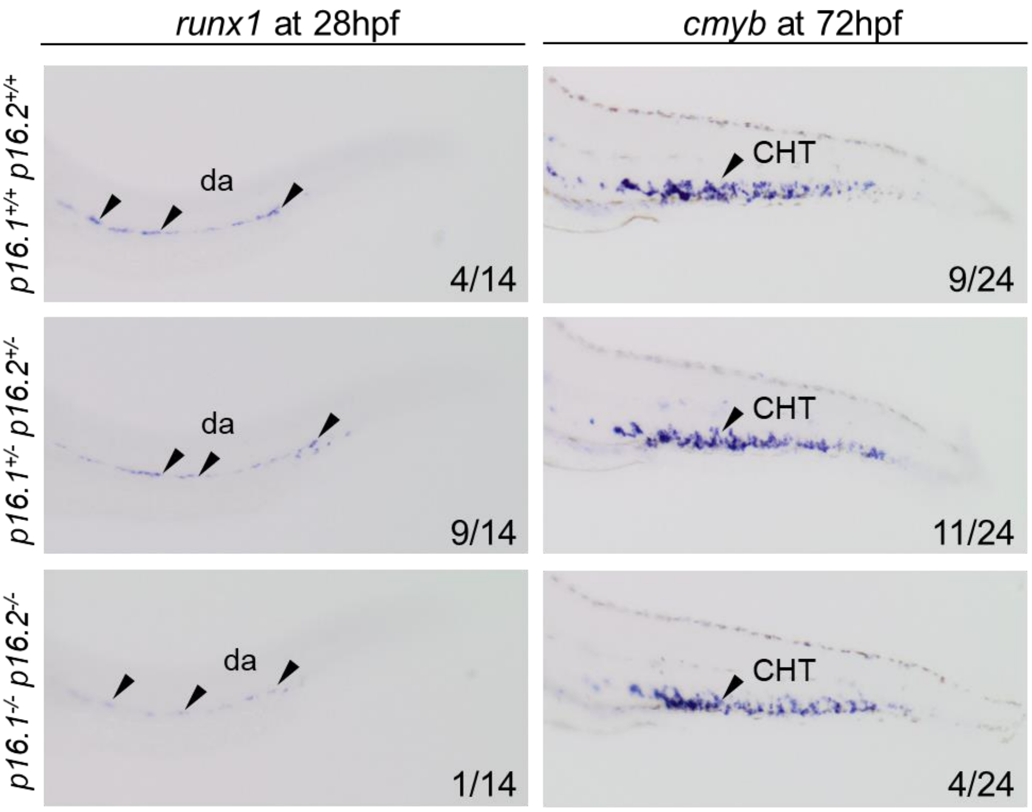


**Supplemental figure S7. Double mutant embryos present normal *runx1* and *cmyb* signal by WISH as compared with wild type siblings.** WISH against *runx1* and *cmyb* at 28hpf and 72hpf, respectively, in the progeny of *p16.1*$^{+/-}$*p16.2*$^{+/-}$ incrossed shows that double knock-out embryos (*p16.1*$^{-/-}$ *p16.2*$^{-/-}$) present normal *runx1* and *cmyb* signal as compared with double heterozygous (*p16.1*$^{+/-}$ *p16.2*$^{+/-}$) and wild type siblings (*p16.1*$^{+/+}$ *p16.2*$^{+/+}$). The embryos were genotyped after imaging.

**Supplemental Figure S8**

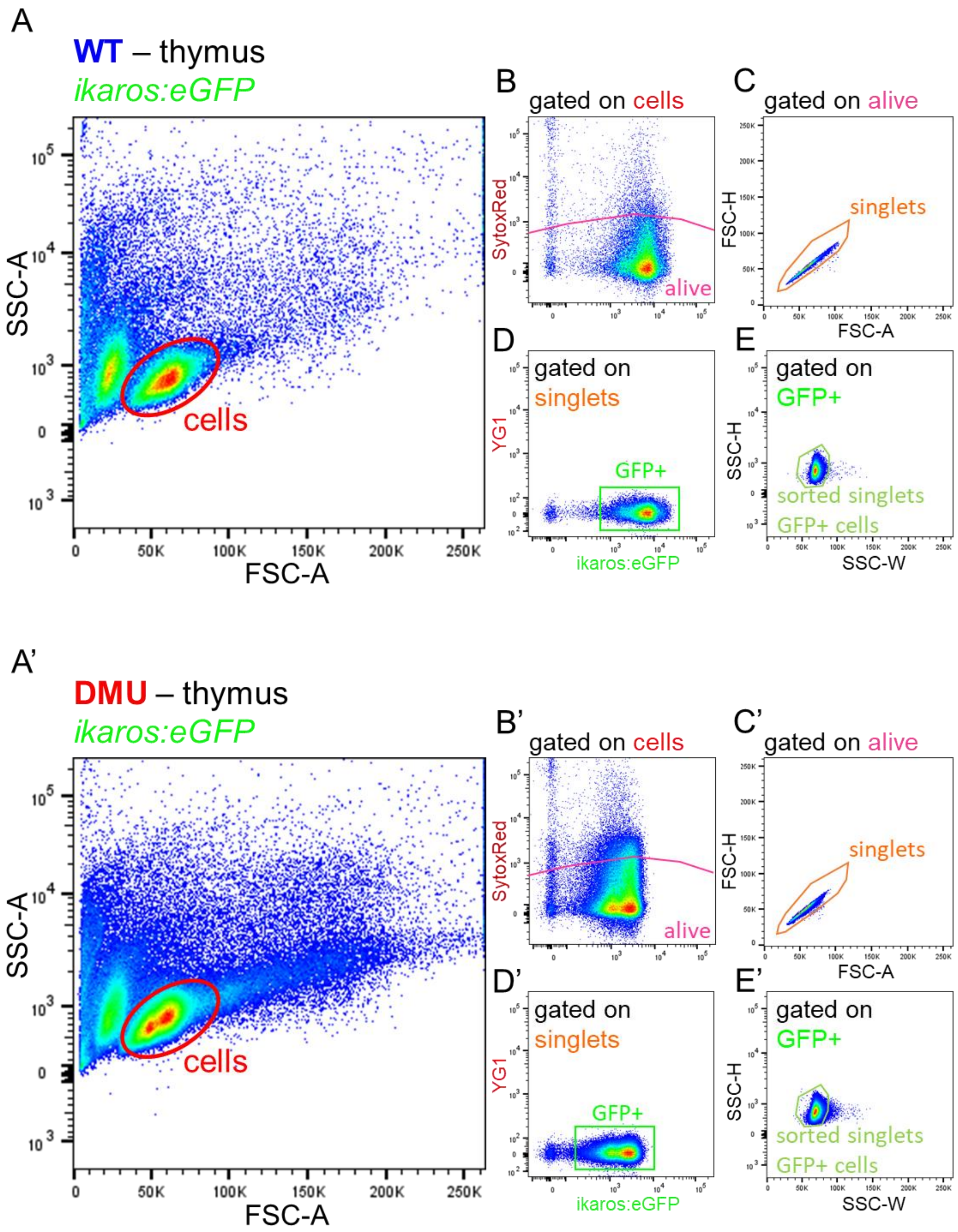

**Supplemental figure S8. Gating strategy to isolate adult thymocytes from the thymus of WT and DMU fish by flow cytometry.** (A and A') Gating on lymphoid cells (cells) within the thymus using FSC/SSC parameters. (B and B') Selection of living SytoxRed$^{-}$ cells (alive) within lymphoid fraction. (C and C') First double exclusion within living cells (alive) by selecting singlets. (D and D') Selection of thymocytes (GFP+). (E and E') Second double exclusion within thymocytes (GFP+) by selecting singlets.

**Supplemental Figure S9**

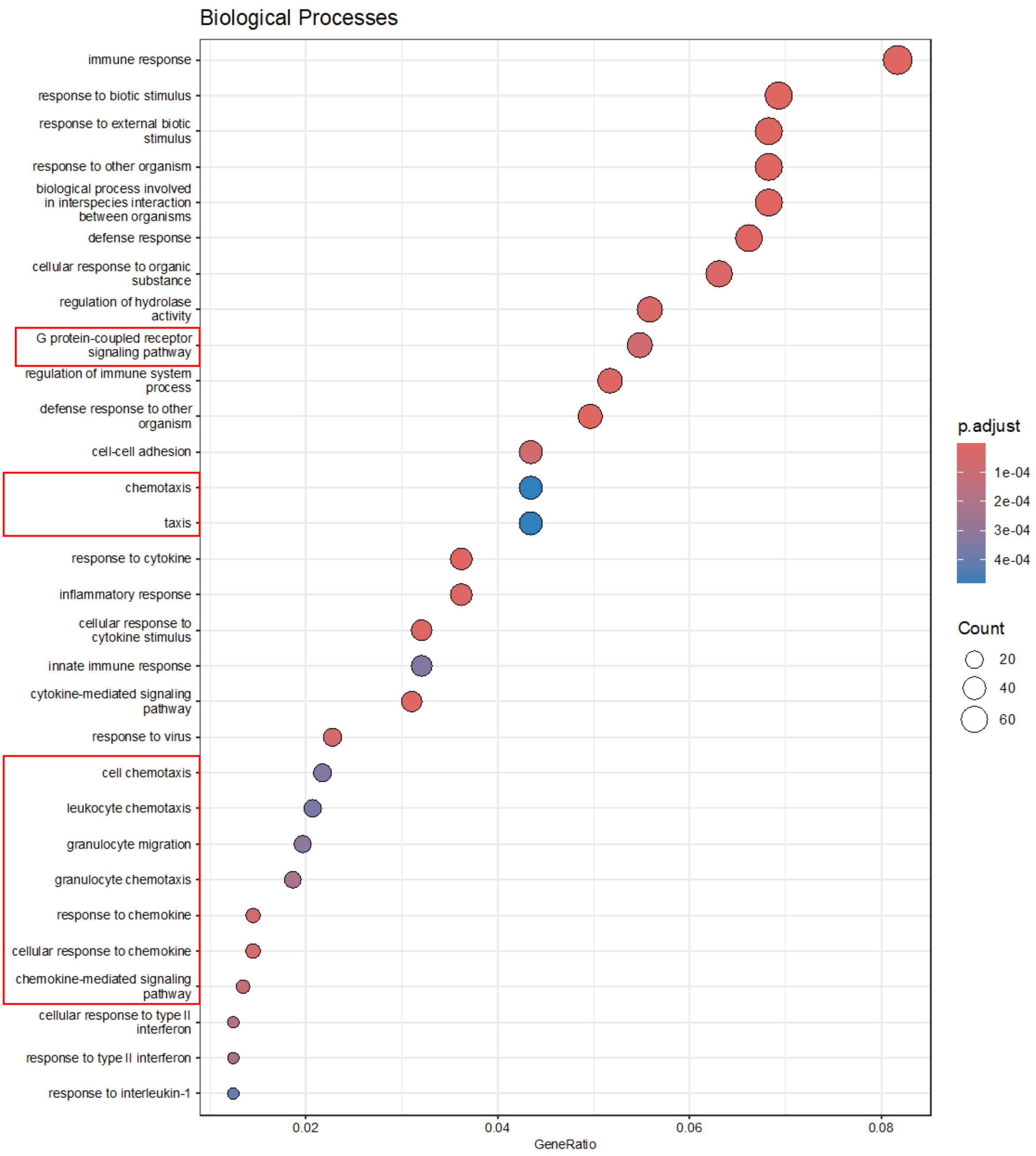


**Supplemental figure S9. GO term analysis performed on differentially expressed genes between WT and DMU thymocytes.** Genes differentially expressed in WT and DMU thymocytes are associated with GO terms related to the G protein-couple receptors signaling pathway, chemotaxis and cell migration.
